# Covalent targeting as a common mechanism for inhibiting NLRP3 inflammasome assembly

**DOI:** 10.1101/2023.06.01.543248

**Authors:** Caroline Stanton, Jie Sun, Kayla Nutsch, Jessica D. Rosarda, Thu Nguyen, Chloris Li-Ma, Sergei Kutseikin, Enrique Saez, John R. Teijaro, R. Luke Wiseman, Michael J. Bollong

## Abstract

The NLRP3 inflammasome is a cytosolic protein complex important for the regulation and secretion of inflammatory cytokines including IL-1β and IL-18. Aberrant overactivation of NLRP3 is implicated in numerous inflammatory disorders. However, the activation and regulation of NLRP3 inflammasome signaling remains poorly understood, limiting our ability to develop pharmacologic approaches to target this important inflammatory complex. Here, we developed and implemented a high-throughput screen to identify compounds that inhibit inflammasome assembly and activity. From this screen we identify and profile inflammasome inhibition of 20 new covalent compounds across 9 different chemical scaffolds, as well as many known inflammasome covalent inhibitors. Intriguingly, our results indicate that NLRP3 possesses numerous reactive cysteines on multiple domains whose covalent targeting blocks activation of this inflammatory complex. Specifically, focusing on compound VLX1570, which possesses multiple electrophilic moieties, we demonstrate that this compound allows covalent, intermolecular crosslinking of NLRP3 cysteines to inhibit inflammasome assembly. Our results, along with the recent identification of numerous covalent molecules that inhibit NLRP3 inflammasome activation, suggests that NLRP3 serves as a cellular electrophile sensor important for coordinating inflammatory signaling in response to redox stress. Further, our results support the potential for covalent cysteine modification of NLRP3 for regulating inflammasome activation and activity.

## Introduction

The NLRP3 (NOD-, LRR- and pyrin domain-containing protein 3) inflammasome is a cytosolic protein complex regulated by the pattern recognition receptor NLRP3 that controls the processing and secretion of multiple pro-inflammatory cytokines including IL-1β and IL-18. While other types of inflammasomes respond to specific toxins or bacterial signals, the NLRP3 inflammasome responds to a broad range of pathogen and danger associated molecular patterns and is therefore most directly associated with sterile inflammation implicated in disease. Activation of the NLRP3 inflammasome requires two signals for assembly and activity. The first signal, priming, is achieved through induction of NFκB which upregulates expression of inflammasome components including NLRP3 and pro-inflammatory cytokines [1, 2]. The second signal, activation, can be achieved through several mechanisms; the most common signal is K^+^ efflux, although Ca^2+^ flux, mitochondrial reactive oxygen species, lysosomal dysfunction, and metabolic changes have also been proposed to activate this complex [3-9]. In response to both signals, NLRP3 oligomerizes and recruits other inflammasome components including NEK7 (NIMA-Related Kinase 7), ASC (Apoptosis-associated speck-like protein containing a caspase recruitment domain), and pro-caspase-1 that assemble into the inflammasome [10, 11]. Inflammasome assembly precipitates the autoproteolysis of pro-caspase 1 to active caspase 1, which then cleaves the pro-inflammatory cytokines, pro-IL-1β and pro-IL-18, to their active forms for cellular secretion [12]. As mature IL-1β and IL-18 are potent inflammatory signaling molecules, NLRP3 inflammasome activity must be tightly regulated to ensure it can properly respond to cellular threats without inducing inflammation that contributes to disease pathology.

One disease most directly associated with NLRP3 inflammasome overactivity is cryopyrin associated periodic syndromes (CAPS), which is caused by hyperactive NLRP3 mutants [13, 14]. Furthermore, overactivation of the NLRP3 inflammasome and excessive pro-inflammatory cytokine production has been implicated in the pathogenesis of numerous inflammatory diseases including gout, rheumatoid arthritis, diabetes, multiple sclerosis, and cardiovascular disease [15-23]. Intriguingly, pharmacologic inhibition or genetic deficiencies of NLRP3 in rodent models of these various inflammatory diseases has been shown to be beneficial. Thus, there has been significant interest in the development of NLRP3 inhibitors as potential therapeutics [24-26]. Despite this potential, clinically approved NLRP3 inflammasome inhibitors have not yet been established, and there is still much yet to be defined regarding the regulation and activation of the NLRP3 inflammasome.

Currently, there are numerous NLRP3 inhibitors in development, many of which bind in the ATP binding pocket of NLRP3. One NLRP3 inhibitor, the sulfonylurea MCC950, has shown positive results in several animal models of disease, yet failed to progress from initial human clinical trials, due to increased liver toxicity [27, 28]. Another inhibitor, OLT1177, thought to block the ATPase activity of NLRP3, completed phase II clinical trials for gout and osteoarthritis, highlighting the potential for targeting NLRP3 through the ATP binding pocket [29]. Post-translational modification such as phosphorylation and ubiquitination also regulate NLRP3 activity [30-34]; however, more recently covalent modification of NLRP3 cysteines by electrophilic chemicals has emerged as a common route for modulating the activity of this complex. Numerous new NLRP3 inhibitors have taken advantage of this cysteine-dependent regulation. Oridonin, a known anti-inflammatory agent, was suggested to inhibit NLRP3 inflammasome activity through modification of C279 of NLRP3 [35]. Another covalent inhibitor of NLRP3, RRx-001, an anticancer agent, modifies C409 of NLRP3 [36]. The immunomodulatory metabolite, itaconate, inhibits NLRP3 activity in mice by modifying C548 of NLRP3 [37]. All of these compounds inhibit the interaction between NLRP3 and NEK7, preventing inflammasome assembly and activation. Other compounds including Bay-11-7082 and MNS are also predicted to covalently modify cysteines in the NACHT domain of NLRP3 but inhibit ATPase activity rather than NEK7 interactions [38, 39].

Further interrogation of covalent NLRP3 inhibition may provide an alternative avenue to learn more about the mechanism of NLRP3 activation and identify new opportunities for the future development of NLRP3 inflammasome inhibitors. Here, we employed a high-throughput screen (HTS) of diverse chemical libraries to identify >450 new inhibitors of NLRP3 inflammasome assembly across numerous chemical scaffolds. Intriguingly, the vast majority of these compounds appear to be covalent modifiers of NLRP3 cysteines, suggesting that NLRP3 is highly sensitive to electrophilic modification. We demonstrate that these covalent compounds, including known NLRP3 inhibitors, modify NLRP3 on multiple cysteines in various domains. These results indicate that NLRP3 is likely a highly sensitive electrophile sensor within the cell, revealing insights into the regulation of this complex during conditions of stress and further motivating the development of new covalent ligands to inhibit NLRP3 inflammasome activity in the context of disease.

## RESULTS AND DISCUSSION

### HTS Identification of inhibitors of NLRP3 inflammasome assembly and activity

To identify novel inhibitors of NLRP3 inflammasome activation, we developed a high-content, imaging-based screening assay of inflammasome assembly using THP1-ASC-GFP cells (Invivogen). These cells express GFP-tagged ASC under control of a NFκB-regulated promoter. In the absence of stimuli, ASC-GFP is diffuse throughout the cell; however, upon administration of an activating stimulus such as ATP or nigericin, ASC-GFP forms discrete puncta, referred to as ASC specks, which reflects inflammasome assembly (**Fig. 1a,b, Fig. S1a**). We monitored ASC speck formation in THP1-ASC-GFP cells in 384-well format through induction of specks in cells pretreated with LPS (priming signal), followed by treatment with nigericin (activation signal) (**Fig. 1a**). We then quantified specks per cell by defining the number of GFP puncta in individual cells. Treatment with nigericin significantly increased ASC specks in THP1-ASC-GFP cells primed with LPS (**Fig. 1c**). Pre-treatment with known inflammasome inhibitors such as MCC950 or oridonin robustly inhibited speck formation (Z’ = 0.8 for MCC950), confirming the sufficiency of this assay for HTS.

**Figure 1:**
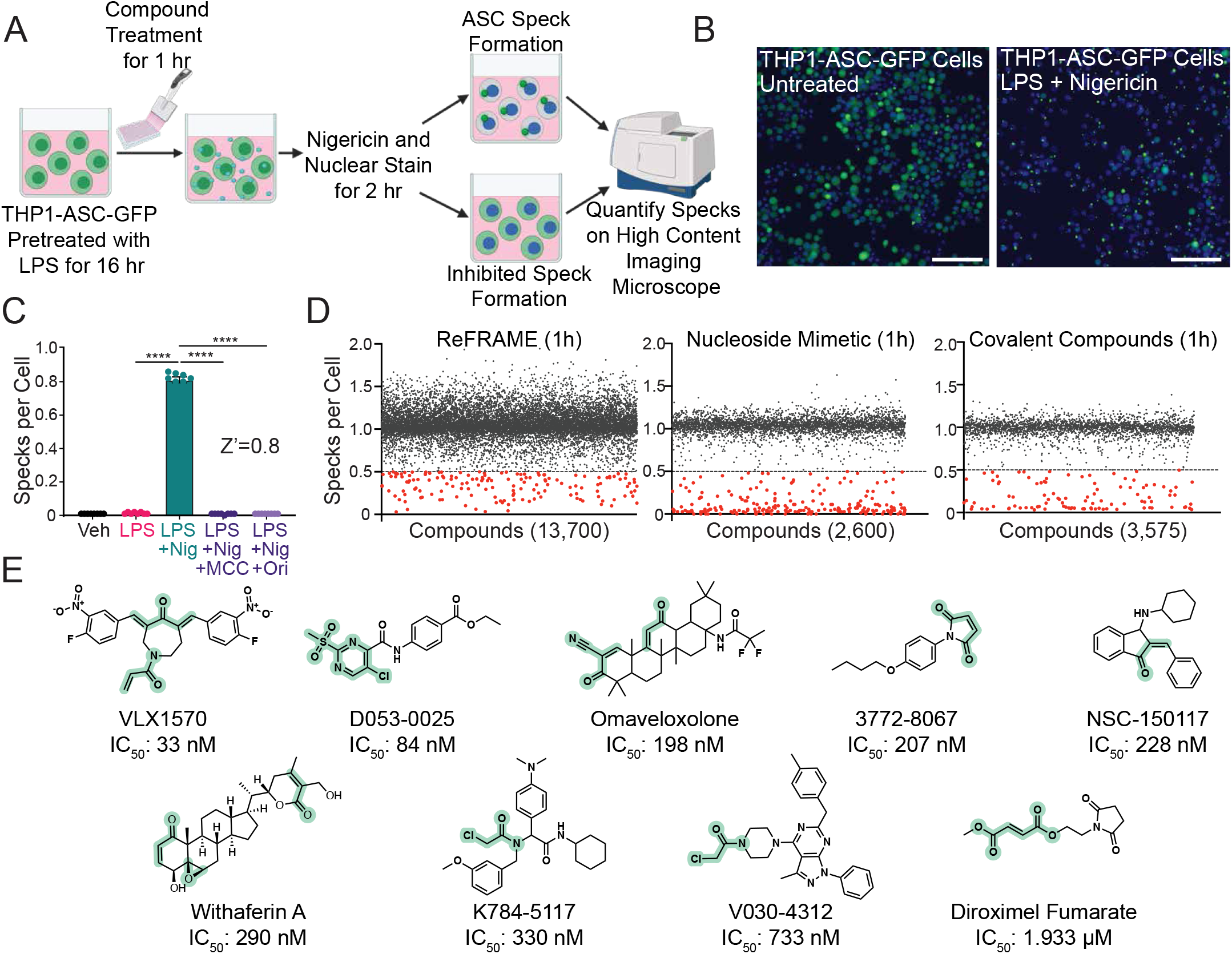
A high content imaging-based screen identifies covalent inhibitors of NLRP3 inflammasome assembly. (A) Schematic of a high throughput screening assay for identifying inhibitors of inflammasome assembly in THP1-ASC-GFP cells primed with LPS, pretreated with compound, and stimulated with Nigericin. (B) Representative images of ASC-GFP from untreated THP1-ASC-GFP cells or ASC-Specks from cells treated with LPS and Nigericin, as depicted in A. Scale bar = 100 μM. (C) Specks per cell of THP1-ASC-GFP cells primed overnight with LPS (1 μg/mL), pretreated with MCC950 (10 μM) or Oridonin (10 μM) for 1 hr, and then stimulated with Nigericin (10 μM) for 2 hr. Error bars show SEM for n = 8 replicates. \*\*\*\**P*<0.0001 for ordinary one-way analysis of variance (ANOVA) with Tukey correction for multiple comparisons between conditions. (D) Scatterplots of specks per cell from each screening well from high throughput screens of the ReFRAME library, Nucleoside Mimetic library, and a Covalent Inhibitor Library. (E) Chemical structures of 9 representative NLRP3 assembly inhibitor scaffolds and their inhibitory potencies in the THP1-ASC-GFP speck assay. Putative covalent reactive groups are depicted in green.

We then used this imaging-based assay to screen the Calibr ReFRAME library, comprised of ∼14k compounds that have undergone extensive preclinical or clinical testing, for inhibitors of NLRP3 inflammasome assembly [40]. Initially, we performed this screen by pre-treating cells for 24 h with compound prior to nigericin administration. This identified 111 hits that inhibited inflammasome assembly (**Fig. S1b**), which included 17 inhibitors of HSP90, a known mechanism of NLRP3 inflammasome inhibition [41, 42]. Intriguingly, we also identified 15 compounds with known cysteine reactive moieties, suggesting these compounds inhibit inflammasome assembly through a covalent mechanism. To better define this mechanism, we rescreened the ReFRAME library in THP1-ASC-GFP cells pre-treated with compounds for 1 h – a timepoint that would minimize potential off-target inhibitory mechanisms and increase identification of direct inhibitors of inflammasome assembly. We also expanded this screen to include a library of covalent compounds and a library of nucleoside mimetics. This more focused screen identified 487 compounds that inhibited inflammasome assembly (**Fig. 1d**). While the libraries screened were chemically diverse and contained many non-covalent scaffolds, an overwhelming number of top hits contained electrophilic moieties suggesting they likely act by a covalent mechanism (**Table 1**). From this screen, we selected 20 of the most potent compounds from 9 potentially covalent, structurally diverse scaffolds for further characterization (**Fig. 1e, Fig. S2a**). We showed that these compounds inhibit inflammasome assembly with IC_50_ values ranging from 30 nM to 2 μM (**Table 1**).

**Table 1:** Inflammasome Inhibition Values for Covalent Inhibitors Identified from High Throughput Screen IC_50_ values for ASC-GFP speck inhibition in THP1-ASC-GFP cells, IL-1β secretion inhibition in WT THP-1 cells when stimulated with LPS and ATP, and protection in WT THP-1 cells from LPS and Nigericin-induced pyroptotic cell death. Maximum protection given relative to primed THP-1 cells not treated with Nigericin (100% viable). LD_50_ for THP-1 cells primed overnight with 1 ng/mL LPS and treated with compound for 24 hr.

| Compound ID | Speck IC<sub>50</sub><br>(μM) | IL-1β IC<sub>50</sub><br>(μM) | Max Protection<br>(%) | Protection IC<sub>50</sub><br>(μM) | 24hr LD<sub>50</sub><br>(μM) |
| --- | --- | --- | --- | --- | --- |
| VLX1570 | 0.033 | 0.391 | 60.7 | 0.072 | 0.293 |
| D053-0367 | 0.070 | 0.273 | 67.05 | 0.131 | 0.766 |
| D053-0156 | 0.0823 | 0.249 | 58.2 | 0.120 | 0.632 |
| D053-0025 | 0.0844 | 1.330 | 53.9 | 0.384 | 1.291 |
| D053-0085 | 0.0854 | 0.368 | 43.45 | 0.110 | 0.574 |
| D053-0257 | 0.0853 | 2.120 | 54.4 | 0.241 | 1.925 |
| D053-0171 | 0.0919 | 0.196 | 44.6 | 0.119 | 0.350 |
| Omaveloxolone | 0.198 | - | 50.9 | 0.167 | 0.145 |
| NSC-150117 | 0.228 | 0.923 | 55.9 | 0.232 | 0.801 |
| 3772-8067 | 0.207 | 6.637 | 34.5 | >20 | 2.755 |
| 4366-0278 | 0.324 | 5.743 | 29.8 | >20 | 2.364 |
| Withaferin A | 0.290 | 0.208 | 63.4 | 0.258 | 0.364 |
| K784-5117 | 0.330 | 0.630 | 53.78 | 0.325 | 12.95 |
| K784-3647 | 0.447 | 0.266 | 49.33 | 0.532 | 6.679 |
| K784-5693 | 0.450 | 0.881 | 44.93 | 0.458 | >20 |
| K784-3648 | 0.587 | 0.513 | 50.08 | 0.656 | 12.41 |
| K784-3666 | 0.842 | 1.513 | 41.05 | 6.183 | 22.83 |
| K784-3025 | 1.479 | 0.790 | 46.68 | 2.308 | 7.202 |
| V030-4312 | 0.733 | - | - | - | - |
| Diroximel Fumarate | 1.933 | - | 71.3 | 1.945 | 16.8 |

Given that inflammasome activation leads to the secretion of the pro-inflammatory cytokine IL-1β and pyroptotic cell death, we established secondary screening assays to evaluate if the identified inhibitors of inflammasome assembly also decreased IL-1β secretion from WT THP-1 cells co-treated with LPS and ATP (HEK-Blue IL-1β reporter assay) and reduced pyroptotic cell death of WT THP-1 cells treated with LPS and nigericin (measured by CellTiter-Glo). We confirmed that pre-treatment with the established NLRP3 inhibitor MCC950 inhibited IL-1β secretion and prevented inflammasome-dependent cell death of WT THP-1 cells (**Fig. S2b,c**). Similarly, all prioritized NLRP3 inhibitors identified in our HTS decreased IL-1β secretion and improved viability of stimulated WT THP-1 cells (**Table 1**). These results confirm that, apart from inflammasome assembly, NLRP3 inhibitors block downstream activities of this complex implicated in inflammatory disease.

### VLX1570 promotes NLRP3 crosslinking

Prioritized inhibitors identified in our HTS all contain potential sites of covalent modification that indicate these compounds likely act through a covalent mechanism. To initially test if these compounds act covalently, we monitored ASC speck formation in stimulated THP1-ASC-GFP cells pretreated with glutathione (GSH) prior to the addition of covalent inhibitor. We found that IC_50_ for speck formation increased ∼10-fold for compounds administered in the presence of GSH (**Fig. 2a**), consistent with a covalent targeting mechanism. We also evaluated the toxicity of these compounds by measuring changes in viability of WT THP-1 cells 24 h after treatment. We observed some toxicity for select compounds at higher doses (**Table 1**); however, this toxicity did not preclude us from utilizing these compounds for monitoring NLRP3 inhibition at shorter timepoints.

**Figure 2:**
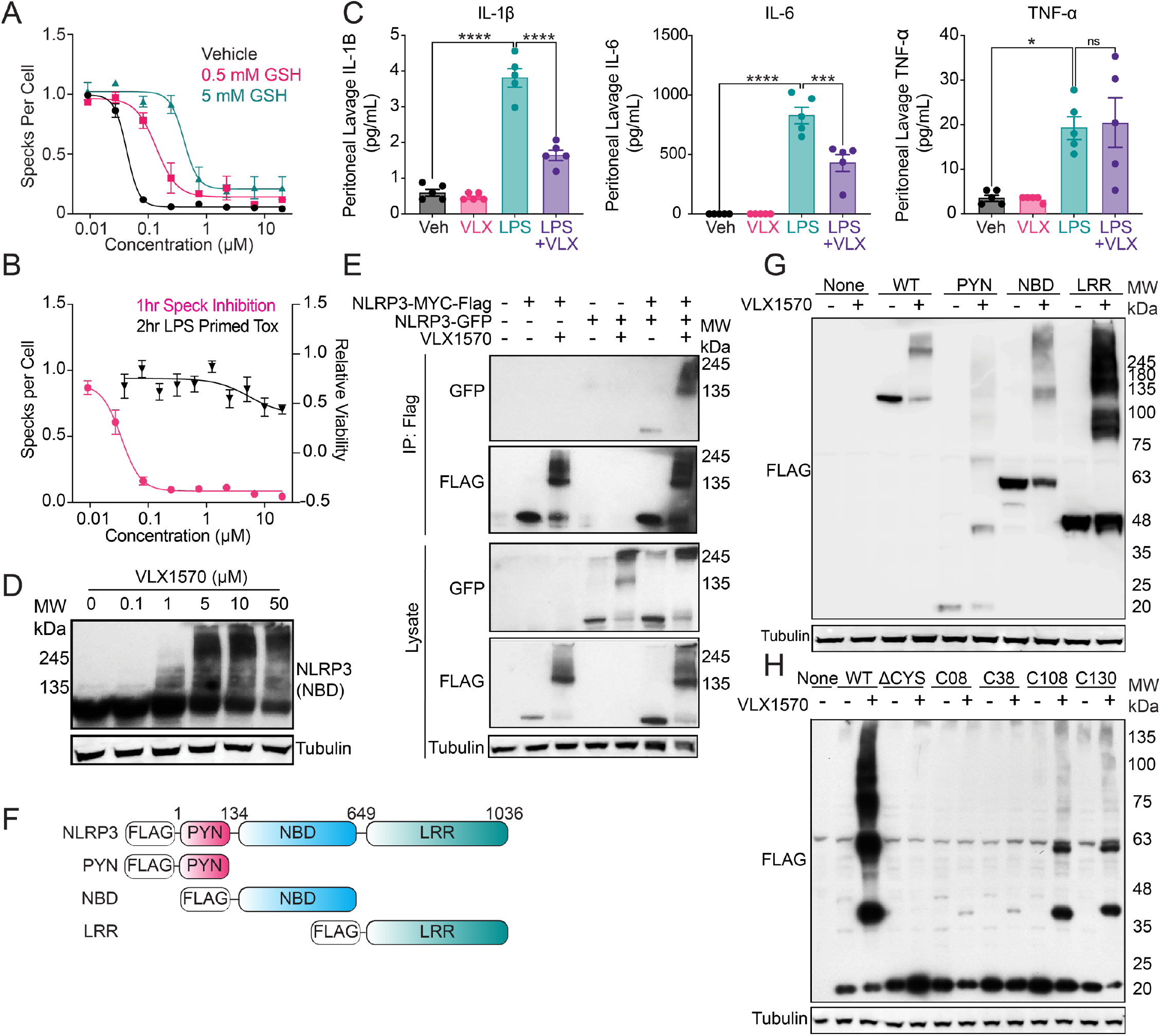
VLX1570 promotes the intermolecular crosslinking of NLRP3 via covalent modification of several cysteines. (A) Representative dose response curves for 1 hr speck inhibition by VLX1570 in THP1-ASC-GFP cells pre-treated with 0, 0.5 or 5 mM GSH. Error bars show SEM for n = 3 replicates. (B) Dose response curves for 1 hr speck inhibition in THP1-ASC-GFP and 2 hr cell viability in THP-1 cells treated with VLX1570. Error bars show SEM for n = 10 replicates (Specks), n = 4 (Viability). (C) IL-1β, IL-6, and TNF-α cytokine levels in the peritoneal lavage fluid from mice pretreated with 4.4 mg/kg VLX1570 (1 hr) and then stimulated with 10 mg/kg LPS (3 hr) evaluated with the Bio-Plex Pro Mouse Cytokine Immunoassay. Error bars show SEM for n = 5 replicates. (D) Western blot of NLRP3 (NBD) and Tubulin from WT THP-1 cells treated in dose response with VLX1570 for 2 hr. (E) Western blot for GFP and FLAG from anti-FLAG immunoprecipitated material from HEK239T cells overexpressing NLRP3-GFP and NLRP3-MYC-FLAG treated with 10 μM VLX1570 (1 hr). (F) NLRP3 Domain constructs. (G) Western blot for FLAG and Tubulin in HEK293T cells overexpressing the indicated FLAG-Tagged NLRP3 domain treated with 10 μM VLX1570 for 2 hr. (H) Western blot of FLAG from HEK293T cells overexpressing NLRP3-FLAG PYN domain constructs with cysteines mutated and individually reintroduced, treated with 10 μM VLX1570 (1 hr).

The most potent compound identified in our screen was VLX1570 (**Fig. 1e, Table 1, Fig. 2b**) – a compound that was developed as a covalent inhibitor of deubiquitinating enzymes (DUBs) including USP14 and UCHL5 with an IC_50_ of DUB inhibition of ∼10 μM *in vitro* [43, 44]. Thus, we sought to better define the activity of this compound. Initially, we monitored the *in vivo* activity of VLX1570 to inhibit inflammasome-associated pro-inflammatory signaling. Towards that aim, we IP administered VLX1570 1 h prior to IP administration of LPS. We then collected peritoneal fluid and monitored levels of pro-inflammatory cytokines 3 hr after LPS administration. Interestingly, we found that VLX1570 reduced levels of both IL-1β and IL-6 in these animals (**Fig. 2c**). However, VLX1570 did not influence TNF-α – a cytokine that is not regulated by inflammasome activity. We did observe reductions in both pro-IL-1β and NLRP3 in peritoneal cells by immunoblotting (**Fig. S3a**). This suggests decreased immune infiltration in the peritoneum. These results indicate that VLX1570 inhibits inflammasome-dependent pro-inflammatory signaling in LPS-treated mice.

VLX1570 contains three potential sites for covalent modification (**Fig. 1e**). Intriguingly, immunoblotting shows that treatment with VLX1570 increases high molecular weight (HMW) populations of NLRP3 in LPS-treated THP1 cells (**Fig. 2d, Fig. S3b**). This appears specific to NLRP3, as other inflammasome components NEK7, ASC, or CASP1 did not show similar effects (**Fig. S3c**). We observed increased levels of HMW NLRP3 in insoluble fractions from VLX1570-treated THP1 cells, suggesting that this modification reduces protein solubility (**Fig. S3d**). VLX1570-dependent increases in HMW NLRP3 was also observed in HEK293T cells overexpressing NLRP3-MYC-FLAG (**Fig. S3e,f**). Interestingly, NLRP3-GFP efficiently co-eluted with HMW NLRP3-MYC-FLAG in FLAG IPs from VLX1570-treated HEK293T cells co-overexpressing these proteins, indicating that these HMW species contain multiple crosslinked NLRP3 proteins (**Fig. 2e**). However, we did not observe USP14 or UCHL5 in NLRP3 immunopurifications from THP1 cells treated with VLX1570, indicating that these HMW NLRP3 species do not contain known targets of VLX1570 (**Fig. S3g**). We showed that recombinant NLRP3 also forms VLX1570-dependent HMW species indicating these HMW species are comprised of intermolecularly cross-linked NLRP3 molecules and the formation of which is not dependent on alternative cellular functions or additional proteins **(Fig. S3h)**.

NLRP3 contains 45 cysteine residues localized throughout its pyrin (PYN), nucleotide binding (NBD), and leucine-rich repeat (LRR) domains (**Fig. 2f**). To identify the specific domain responsible for VLX1570-dependent induction of HMW species, we overexpressed these individual domains and monitored formation of HMW species by immunoblotting. Surprisingly, we found that each overexpressed domain showed increased formation of HMW species, indicating that VLX1570 can induce crosslinking across multiple different cysteine residues (**Fig. 2g**). This suggests that VLX1570 specifically crosslinks proteins containing multiple electrophile sensing cysteine residues. Thus, to identify specific cysteines modified by VLX1570, we specifically focused on the NLRP3 pyrin domain, which contains only 4 cysteines (**Fig. S3k**). Importantly, mutation of all cysteines within the pyrin domain to serine completely blocked VLX1570-dependent induction of HMW species (**Fig. 2h**). Next, we re-introduced these cysteines back into the cysteine-less pyrin domain to identify those specifically modified by VLX1570. Interestingly, while we found that all cysteines showed some modification, Cys 108 and Cys 130 were shown to be the two cysteine residues that most strongly induced formation of HMW species (**Fig. 2h**). This indicates that these two cysteines may be more sensitive to electrophilic modification. Recombinant pyrin domain, only retaining cysteine 108, forms HMW pyrin species confirming C108 as a site of modification for VLX1570 **(Fig. S3i,j)**. This further highlights the ability of this compound to modify multiple cysteines on NLRP3. We propose that VLX1570 may target highly reactive cysteines such as those found on electrophile sensors. Consistent with this, we found that VLX1570 also induced crosslinking of KEAP1-FLAG – another electrophile sensor containing multiple reactive cysteine residues – in HEK293T cells (**Fig. S3l**). Collectively, these results suggest that VLX1570 induces crosslinking of NLRP3 through reactive cysteines on multiple different domains to block inflammasome assembly and activation.

### A 2-Sulfonylpyrimidine-Containing Scaffold Covalently Modifies NLRP3

To further characterize the capacity for electrophile sensing by reactive cysteines on NLRP3, we selected two novel covalent scaffolds, a 2-sulfonylpyrimidine containing scaffold (e.g., D053-0025) and a chloroacetamide containing scaffold (e.g., K784-5117), for further study. These compounds inhibit ASC-speck formation with IC_50_ values of 84 and 330 nM, respectively (**Fig. 1e**). We next performed structure activity profiling of each scaffold to identify specific locations within their structures to incorporate click-compatible alkyne moieties (**Fig. 3a, Fig. S4a-c**). These efforts established that the 2-sulfonylpyrimidine scaffold was a more tractable scaffold for the optimization of inhibitory potency, resulting in the identification of D053-0367 (IC_50_ = 70 nM) **(Fig. 3b,c)** which acted covalently in the GSH shift assay (**Fig. 3d)** and inhibited pyroptotic cell death (**Fig. 3e)**, as well as an alkyne derivatized probe compound (P207-9174, IC_50_ = 70 nM). We confirmed that this probe efficiently inhibited ASC speck formation in THP1-ASC-GFP cells to extents similar to that observed for the parent compound (**Fig. 3f**).

**Figure 3:**
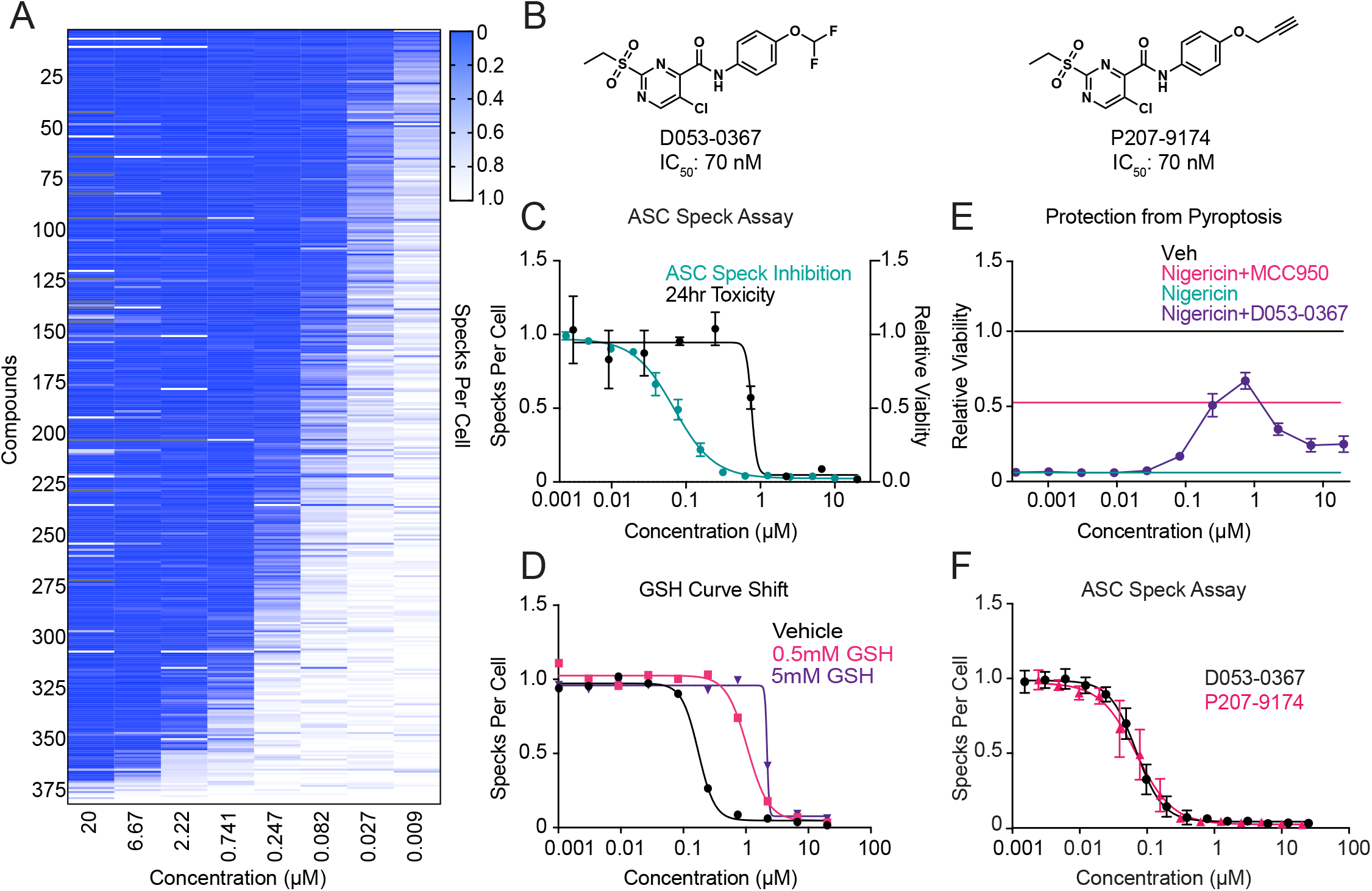
A covalently acting, 2-sulfonylpyrimidine-based inhibitor of NLRP3 assembly. (A) Heat map-based summary of a structure activity relationship depicting relative ASC-Speck assembly from THP1-ASC-GFP cells treated with a library of 2-sulfonylpyrimidines. (B) Structures of D053-0367 and alkyne probe P207-9174. (C) Dose response curves for 1 hr speck inhibition in THP1-ASC-GFP and 24 hr cell viability in THP-1 cells treated with D053-0367. Error bars show SEM for n = 2 replicates. (D) Dose response curves for 1 hr speck inhibition by D053-0367 in THP1-ASC-GFP cells pre-treated with 0, 0.5 or 5 mM GSH. (E) Protection from NLRP3-mediated pyroptotic cell death induced by LPS and Nigericin in WT THP-1 pretreated with D053-0367 (1 hr) in dose response. Error bars show SEM for n = 2 replicates. (F) P207-9174 and D053-0367 inhibition of ASC Specks with 1 hr pretreatment in THP1-ASC-GFP cells. Error bars show SD for n = 6 replicates.

### Covalent inflammasome inhibitors modify cysteines of NLRP3 on multiple domains

The 2-sulfonylpyrmidine scaffold can modify biological thiols covalently, acting through an S_N_Ar based mechanism, resulting in the elimination of the sulfonyl group (**Fig. S5a)**. The nature of this modification was confirmed via mass spectrometry of D053-0367 incubated with GSH (**Fig. S5b)**. We then probed the ability of this compound to covalently modify NLRP3. We treated THP-1 cells with probe compound and then monitored NLRP3 labeling by appending a biotin to the probe using click chemistry followed by streptavidin isolation and immunoblotting for NLRP3. P207-9174 showed dosable modification of NLRP3 using this assay (**Fig. S5c**). Dose dependent modification was also observed for overexpressed FLAG-NLRP3 in HEK293T cells (**Fig. S5d**). Further, we did not observe labeling of other inflammasome subunits such as NEK7 by this probe (**Fig. S5e)**. Importantly, probe-dependent labeling of NLRP3 was inhibited by co-treatment with excess parent compound, demonstrating that this labeling reflected compound modification (**Fig. 4a**). Collectively, these results confirm that the identified 2-sulfonylpyrimidine scaffold covalently modifies NLRP3.

**Figure 4:**
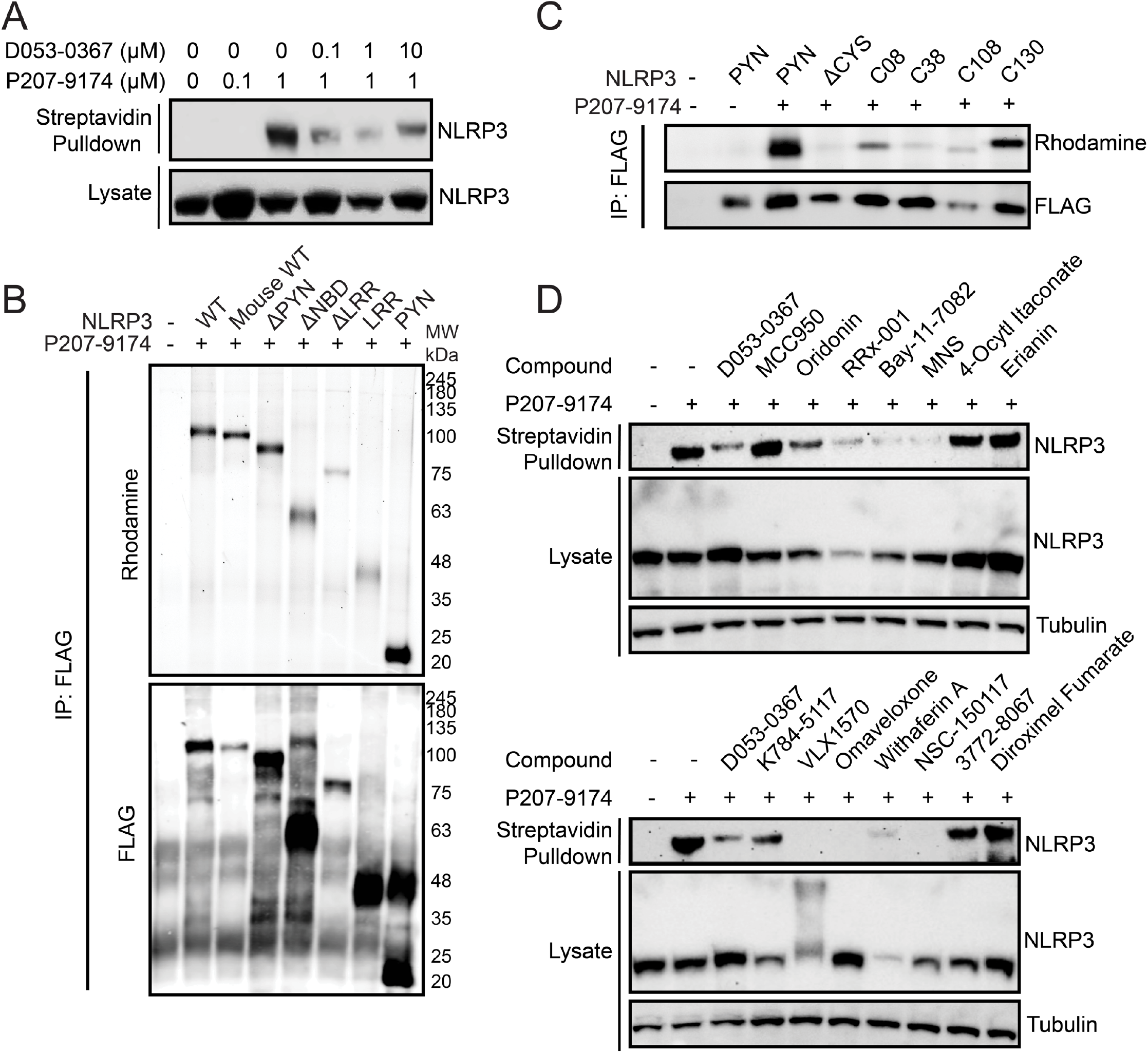
The identified 2-sulfonylpyrimidine scaffold modifies multiple sensor cysteines of NLRP3. (A) Western blot for NLRP3 immunopositivity of streptavidin-enriched material after in situ treatment of THP-1 cells with D053-0367 (1 hr) followed by labeling with P207-9174 (1 hr). (B) Anti-FLAG Western blot and rhodamine imaging of FLAG immunoprecipitated material after in situ treatment of HEK293T cells overexpressing the indicated FLAG-tagged NLRP3 domain treated with 1 μM P207-9174 (1 hr). (C) Anti-FLAG Western blot and rhodamine imaging of FLAG immunoprecipitated material after in situ treatment of HEK293T cells expressing the indicated FLAG-tagged PYN domain cysteine mutants treated with 1 μM P207-9174 (1 hr). (D) Anti-NLRP3 Western blot of streptavidin-enriched material after in situ treatment of THP-1 cells with 1 μM P207-9174 (1 hr) competed with the indicated compounds (10 μM, 1 hr).

We next sought to identify the specific NLRP3 domains modified by the 2-sulfonylpyrimidine probe. We overexpressed FLAG-tagged constructs of individual NLRP3 domains in HEK293T cells and monitored compound modification in FLAG immunopurifications followed by click chemistry-dependent appendage of a fluorescent rhodamine to the probe. Intriguingly, we found that the probe labeled all individually expressed domains of NLRP3, indicating that this compound targets multiple cysteines throughout the protein sequence (**Fig. 4b**). To identify specific cysteines modified by this probe, we again focused on the NLRP3 pyrin domain, which contains only 4 cysteines (**Fig. S3k**). Mutation of all cysteines within the pyrin domain to serine completely blocked probe-dependent labeling (**Fig. 4c**) When these cysteines were re-introduced back into the cysteine-less pyrin domain to identify those specifically modified by our probe, we found that while all cysteines showed some modification, Cys 8 and Cys 130 were shown to be the two cysteine residues most prominently labeled by our probe (**Fig. 4c**). This indicates that these two cysteines may be more sensitive to electrophilic modification. This further highlights the ability of these compounds to modify multiple cysteines on NLRP3.

Numerous established covalent NLRP3 inflammasome inhibitors have been previously suggested to work by targeting specific cysteines on the protein structure. Thus, the ability for our probe to broadly modify multiple cysteines on different domains provides an opportunity to evaluate the specific or broad nature of cysteine targeting by these established compounds and other covalent compounds identified in our screen using a competition assay. Surprisingly, we found that the vast majority of covalent compounds, including the established NLRP3 inhibitors oridonin and RRx-001, competed with our probe in labeling of endogenous NLRP3 in THP1 cells (**Fig. 4d**). In contrast, no competition was observed for non-covalent NLRP3 inhibitors such as MCC950. These results indicate that covalent compounds, including the established inhibitor oridonin and many identified through our screens, appear to modify multiple cysteine residues throughout the NLRP3 sequence, indicating that the protein is highly sensitive to regulation through covalent modification.

## Concluding Remarks

Our data demonstrates that NLRP3 inflammasome assembly can be inhibited by a variety of previously unpublished covalent chemical scaffolds at multiple cysteines. Identification of structurally diverse covalent inhibitors of NLRP3 indicates that NLRP3 may be a good covalent drug target for the treatment of inflammatory diseases through modification of cysteines resulting in inhibitory mechanisms not previously shown. This may be advantageous compared to an ATPase inhibitor, as covalent drugs often display increased specificity, potency, and duration of engagement [45]. Identified here, the deubiquitinase inhibitor VLX1570 inhibits the NLRP3 inflammasome through induction of high molecular weight NLRP3 aggregates via covalent cross-linking. This demonstrates a novel, alternative mechanism for covalent NLRP3 inflammasome inhibition, independent of ATPase activity or NEK7 interactions. We also show that VLX1570 is active *in vivo* and can inhibit NLRP3 inflammasome-dependent inflammation induced by LPS challenge.

In conjunction with previous publications showing covalent modification and inhibition of NLRP3, this data indicates that numerous cysteines in NLRP3, including those in the pyrin domain which have not been previously shown to be modified, may be targeted by covalent molecules of varying chemical scaffolds. However, our data demonstrates the identified covalent inhibitor, VLX1570, the 2-sulfonylpyrimidine scaffold, and previously published inhibitors, are likely not modifying a single cysteine, but an array of NLRP3 cysteines which contribute to the inhibition. Additionally, this modification of multiple cysteines by each compound makes it difficult to increase potency of covalent compounds through structure activity relationship analysis as there is not a single binding site to optimize compound binding. Despite this, the high sensitivity of NLRP3 to covalent modification indicates that *NLRP3 serves as an electrophile sensor within the cell to regulate inflammatory signaling*, providing justification for further covalent NLRP3 inhibitor development for the treatment of inflammatory disease. With further characterization of covalent inhibition at additional sensor cysteines on NLRP3, more potent and specific NLRP3 inhibitors may be developed in the future.

## Author contributions

C.S., J.R.T, E.S., R.L.W., and M.J.B. designed research; C.S., J.S., K.N., J.R., T.N., C.L., and S.K., performed research; C.S. and K.N. analyzed data; and C.S., R.L.W., and M.J.B. wrote the paper.

## Acknowledgements

This work was supported by the NIH (GM146865 to MJB; DK107604 and AG046495 to RLW). We thank Alan Chu (CALIBR) for providing THP1-ASC-GFP, WT THP-1, and HEK-Blue™ IL-1β cells, Luke Lairson and lab members for assistance with CX5 HCS platform, Mannkyu Hong for assistance with cysteine mass spectrometry, and Kristi Marquardt and Nora Leaf for their technical assistance with the Bio-Plex Pro Cytokine assay.

## Materials and Methods

### Compounds, Antibodies, and Plasmids

The ReFRAME Library was accessed through CALIBR at Scripps Research. The covalent compound and the nucleoside mimetic libraries were purchased from ChemDiv. For follow-up assays, compounds identified from ChemDiv libraries were repurchased through ChemDiv. Ultrapure LPS, E. coli 0111:B4 (Invivogen; Cat. No. tlrl-3pelps) was dissolved in ultrapure water and administered at 1 ng/mL. MCC950 (SelleckChem; Cat. No. S7809) was dissolved in water and administered at 10 μM. Nigericin Sodium Salt (Cayman Chem.; Cat. No. 11437-25) was dissolved in ethanol and administered at 10 μM. All other compounds were dissolved in DMSO and sourced as listed in Supplementary Table 1.

### Cell Culture and High Throughput Screening

THP1-ASC-GFP Reporter cells, WT THP-1 cells, and HEK-Blue™ IL-1β cells were obtained from Invivogen and maintained according to protocols supplied by Invivogen. For high throughput screening, THP1-ASC-GFP Reporter cells were primed overnight with 1 ng/mL LPS. Cells were plated in black, clear-bottom 384 well plates at a density of 20,000 cells/well in 50μL of media. Compounds from Covalent Diversity Library and Nucleoside mimetic library were transferred using an Agilent Bravo outfitted with a pintool head to transfer 100 nL of 10 mM compound to each well (final concentration 20 μM). For the Calibr ReFRAME Library, compounds were prespotted using an Echo Acoustic Liquid Handler instrument (Labcyte) before addition of cells to give a final concentration of 1 μM. Following pre-treatment time (1 to 24 hr), 10 μL of media containing Nigericin and Hoechst were added to each well (final concentration 10 μM Nigericin, 5 μg/mL Hoechst). Two hours after addition of Nigericin, cells were imaged on the Cellinsight CX5 HCS Platform. ASC-Specks were quantified using the SpotDetector function of the Cellinsight High Content Analysis Platform. Confirmation assays and dose responses for ASC-Speck formation were performed in a similar manner. GSH Curve shift assays were performed in a similar manner except 10 μL of GSH (final concentration 5 mM, 1 mM, or 0.5 mM) was added 15 min prior to addition of covalent inhibitors.

### IL-1β Secretion Inhibition Assay

WT THP-1 cells were primed and treated with compound as described in ASC-Speck assay. Following pre-treatment of 1 hr with covalent inhibitors, cells were treated with ATP in 10 μL water, pH 7.4 (Final concentration 5 mM). Cells were allowed to secrete IL-1β overnight. The following day, 10,000 HEK-Blue™ IL-1β cells were plated in 30 μL of media in black, clear bottom 384-well plates, and 10 μL of IL-1β conditioned media was added to each well. Cells were incubated overnight to produce SEAP. The following day, 30 μL of QUANTI-Blue (Invivogen) was added to each well and incubated at 37 °C for 30 min-24 hr and absorbance at 655 nM was measured. IL-1β secretion inhibition by compounds was normalized to the maximal inhibition of MCC950.

### Pyroptotic Cell Death Assay

WT THP-1 cells were primed with LPS, pretreated with covalent inhibitor compound, and activated with Nigericin as previously described in the ASC-Speck Assay. 10 μM MCC950 was used as a control to inhibit pyroptotic cell death, and cells not treated with Nigericin were used as a control for no pyroptotic cell death (maximal viability).

Two hours after addition of Nigericin, 30 μL of CellTiter Glo (Promega, diluted 1:6), was added to each well and luminescence was measured with an Envision plate reader. Compound cytotoxicity was measured using a similar protocol where compound was added to LPS-primed WT THP-1 cells and viability was measured 24 hr following compound treatment.

### Immunoblotting and Immunoprecipitation

For expression of NLRP3 in HEK293T cells, 10^6^ cells were plated on poly-d-lysine coated 6-well plates and transfected with 2 μg of DNA per well and 7 μL of Fugene in 100 μL of Optimem. After 48 hr, the cells were treated with compound as desired. For experiments done with endogenous NLRP3, WT THP-1 cells were primed overnight with 1 ng/mL LPS and treated with compound before collection. Cells were rinsed with PBS and each well harvested in RIPA buffer with protease inhibitor and lysed on ice for >15 min. Insoluble material was separated via centrifugation. Concentration of soluble lysates was measured via absorbance with a nanodrop instrument. Lysate samples loaded typically were between 40-100 μg. For samples used for FLAG immunoprecipitation, lysates were normalized to 1 mg/mL and 20 μL of Magnetic FLAG Beads were added to each sample and incubated at 4°C overnight. For NLRP3 (NBD) immunoprecipitation, lysates were normalized to 1 mg/mL and 20μL of Protein G Magnetic Beads and 2 μL of NLRP3 (NBD) antibody were added to each sample and incubated at 4°C overnight. Beads were immobilized using a magnetic Eppendorf holder and washed 2X with lysis buffer and 1X with DPBS. Beads were either resuspended in loading dye for analysis via western blot, or in 100 μL of DPBS for click chemistry. Click Chemistry Master Mix was comprised of 6 μl of 1.7 Tris((1-benzyl-1H-1,2,3-triazol-4-yl)methyl)amine (Sigma) in 4:1 tBuOH:DMSO solution, 2 μl of 50 mM CuSO4 (Sigma) in water, 2 μl of 5 mM rhodamine-azide in DMSO and 2 μl of 50 mM tris(2-carboxyethyl)phosphine (TCEP) in water. To each sample, 12 μL of click chemistry master mix was added and incubated for 1 hr in the dark. Beads were immobilized using a magnetic Eppendorf holder and washed 2X with DPBS. Beads were resuspended in loading dye for analysis via western blot and boiled for 15 min at 95 °C. Protein material was separated via SDS– polyacrylamide gel electrophoresis (SDS-PAGE) (12, 15, or 17 well gels, 4-20% Bis-tris BOLT gels, Invitrogen), with 1X MOPS Buffer (Invitrogen) at 200 V for 40 min. Proteins were transferred to polyvinylidene difluoride (PVDF) membrane with semi-dry transfer at 20 V for 1 hr, and then the PVDF membrane was blocked with 5% blotting grade milk in TBST (20 mM Tris, 137 mM NaCl, 0.1% Tween 20) for 1 hr. Membranes were incubated with respective primary antibodies overnight at 4°C in 5% blotting grade milk or 5% BSA as indicated in Supplementary Table 2. The next day, membranes were washed 6X over 30 min with TBST, and then incubated in secondary antibody (LI-COR, 1:5000), (HRP, 1:3333) in 5% milk for 1 hr. Blots were washed 10X over 1 hr with TBST and either imaged using LI-COR fluorescence imager or incubated in HRP substrate (West Dura) and developed with autoradiography film.

### NLRP3 Target Confirmation and Competition

10 mL of WT THP-1 cells at 1×10^6^ cells/mL were primed overnight with 1 ng/mL LPS and either pre-treated with compound before probe treatment, or treated only with probe, and then collected. Cells were lysed in 1 mL of DPBS using a tip sonicator. Insoluble material was separated via centrifugation. Samples were diluted to 2 tubes of 0.5 mL of 2 mg/mL of lysate and treated each with 55 μL of Biotin Click master mix comprised of 30 μl of 1.7 Tris((1-benzyl-1H-1,2,3-triazol-4-yl)methyl)amine (Sigma) in 4:1 tBuOH: DMSO solution, 10 μl of 50 mM CuSO4 (Sigma) in water, 10 μl of 5 mM biotin-PEG3-azide in DMSO and 10 μl of 50 mM tris(2-carboxyethyl)phosphine (TCEP) in water, and incubated for 1 hr in the dark. Reaction was stopped with addition of 500μL of ice cold methanol and precipitated proteins were pelleted via centrifugation and rewashed with methanol. Protein pellets were resuspended by sonication in 12 mL of 0.1% SDS in DPBS to which 220 μL of pre-equilibrated streptavidin beads (Thermo Scientific, PI20357) were added. Samples were incubated overnight at room temperature and then was washed twice with 0.1% SDS in DPBS, twice with DPBS, and then twice with water, 10 mL each. Beads were resuspended in 200 μL of loading dye with DTT and boiled at 95 °C for 15 min to elute. Immunoblotting was performed as described above.

### In vivo LPS Challenge

To evaluate in vivo efficacy of VLX1570, 5 9-week-old C57BL/6J male mice per group (4 groups total) were treated with 4.4 mg/kg VLX1570 dissolved in PEG/Chremophore/Tween (50/10/40), diluted 1:10 with saline, or treated with the matching volume of vehicle intraperitoneally (IP). Then, 1 hr after dose of VLX1570 or vehicle, animals were IP injected with LPS (L2880; Sigma Aldrich) at 10 mg/kg or matching volume of PBS and mice were sacrificed after 3 hr using CO_2_. Cold PBS (10 mL) was injected into the peritoneal cavity and peritoneal lavage fluid was collected. The peritoneal lavage fluids were centrifuged at 1,000 xg for 5 min, and normalized supernatants were used for cytokine measurement by Bio-Plex Pro Mouse Cytokine Immunoassay. The cells collected from the peritoneal lavage fluid were lysed, normalized, and immunoblotting for NLRP3 and pro-IL-1β performed. Mouse experiments were approved by and conducted in accordance with the guidelines of The Scripps Research Institute IACUC.

### Mass Spectrometry

Electrophilic compound reactions with GSH were done *in vitro*. A 50 mM reduced glutathione (GSH) stock solution in 0.2 M phosphate buffer pH 7.4 was prepared fresh and kept on ice. Compounds stocks dissolved in DMSO at 10 mM were kept at – 20°C for storage. The GSH stock solution was diluted to 4 mM in 50 μL of 0.2 M phosphate buffer pH 7.4. Electrophilic compounds (1 mM) were added by brief vortexing and then incubated at 37 °C for 1 hour in a water bath. Covalent product formation was evaluated by LC-MS by diluting 5 μL of the reaction mixture into 45 μL of 1:1 CH3CN/H2O + 0.01% TFA and injecting 1 μL on an Agilent 6135 instrument.

### Recombinant PYN Domain Expression

NLRP3 PYN Domain (3-110) with C08S, C38S mutations, was cloned into a PET28b vector and transformed into BL21 (DE3) competent E. coli. Starter cultures were grown overnight, and 250 mL cultures inoculated with 2.5 mL each of starter culture. At an OD_600_ of 0.6-0.8, cultures were induced with 0.1 mM IPTG and grown overnight at 18 °C. Cultures were collected by centrifugation and lysed by sonication in lysis buffer containing 20 mM Hepes (pH 7.5), 150 mM NaCl, and 0.5 mM tris(2-carboxyethyl)phosphine (TCEP). Lysate was clarified by centrifugation and soluble lysate was incubated with Ni-NTA Agarose beads (Qiagen, 30210) for 2 hr at 4 °C.

The beads were then added to a column, the unbound fraction removed, and washed with 50 mM imidazole. His-tagged NLRP3 PYN was eluted with 200 mM imidazole and purification was confirmed with SDS-PAGE and Coomassie staining.

---

**Supplementary Figure 1:**
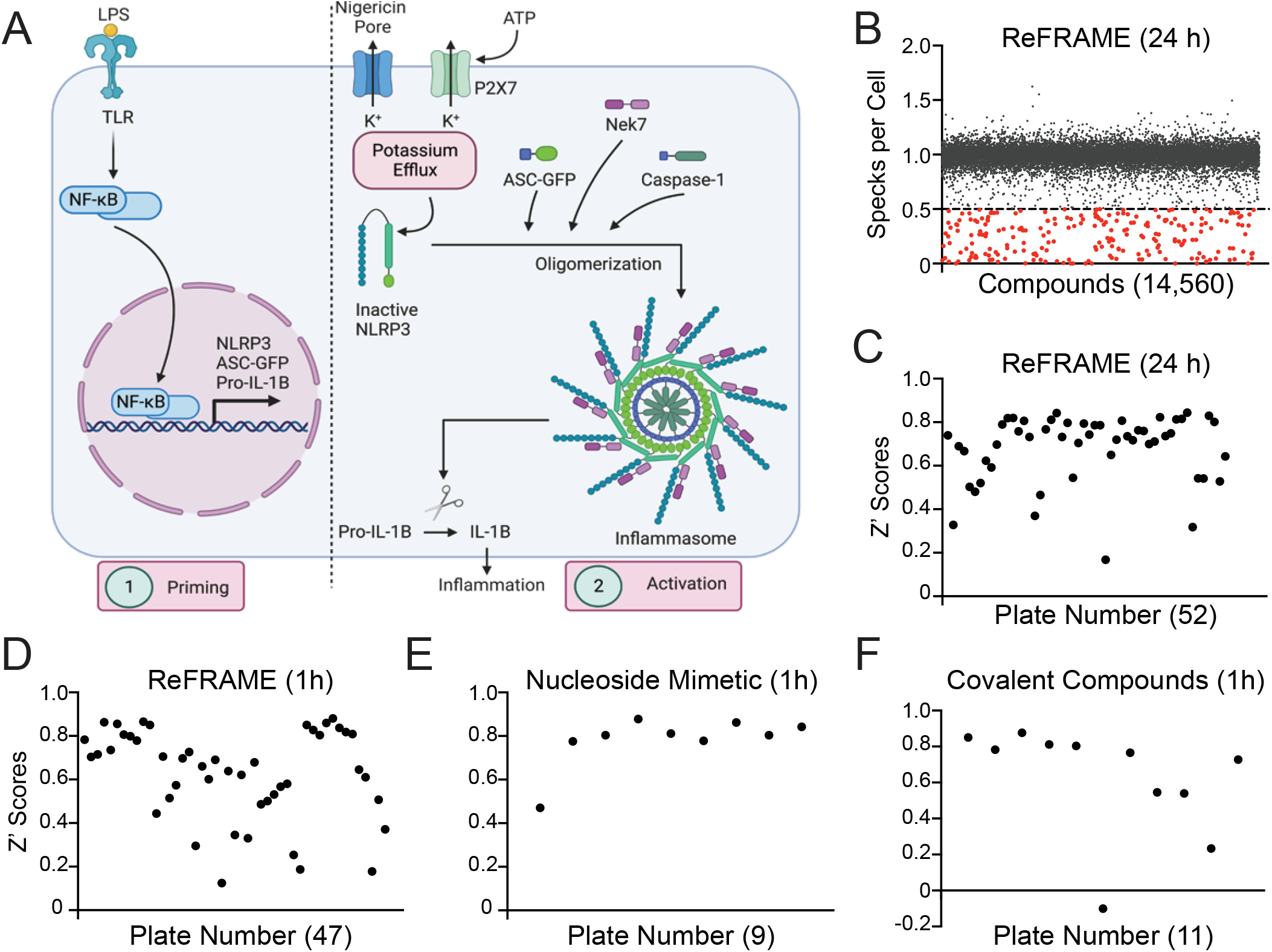
A High Throughput Screen for Inhibitors of ASC-Speck Formation (A) Schematic of NLRP3 inflammasome activation including ASC-GFP. (B) Scatterplots of specks per cell evaluated via high throughput screen for inhibitors of inflammasome assembly for compounds tested in 24 hr ReFRAME Screen. (C-F) Z’ Values for plates from 24 hr ReFrame Screen (C), 1 hr ReFrame Screen (D), 1 hr Nucleoside Mimetic Screen (E), and 1 hr Covalent Library Screen (F).

**Supplementary Figure 2:**
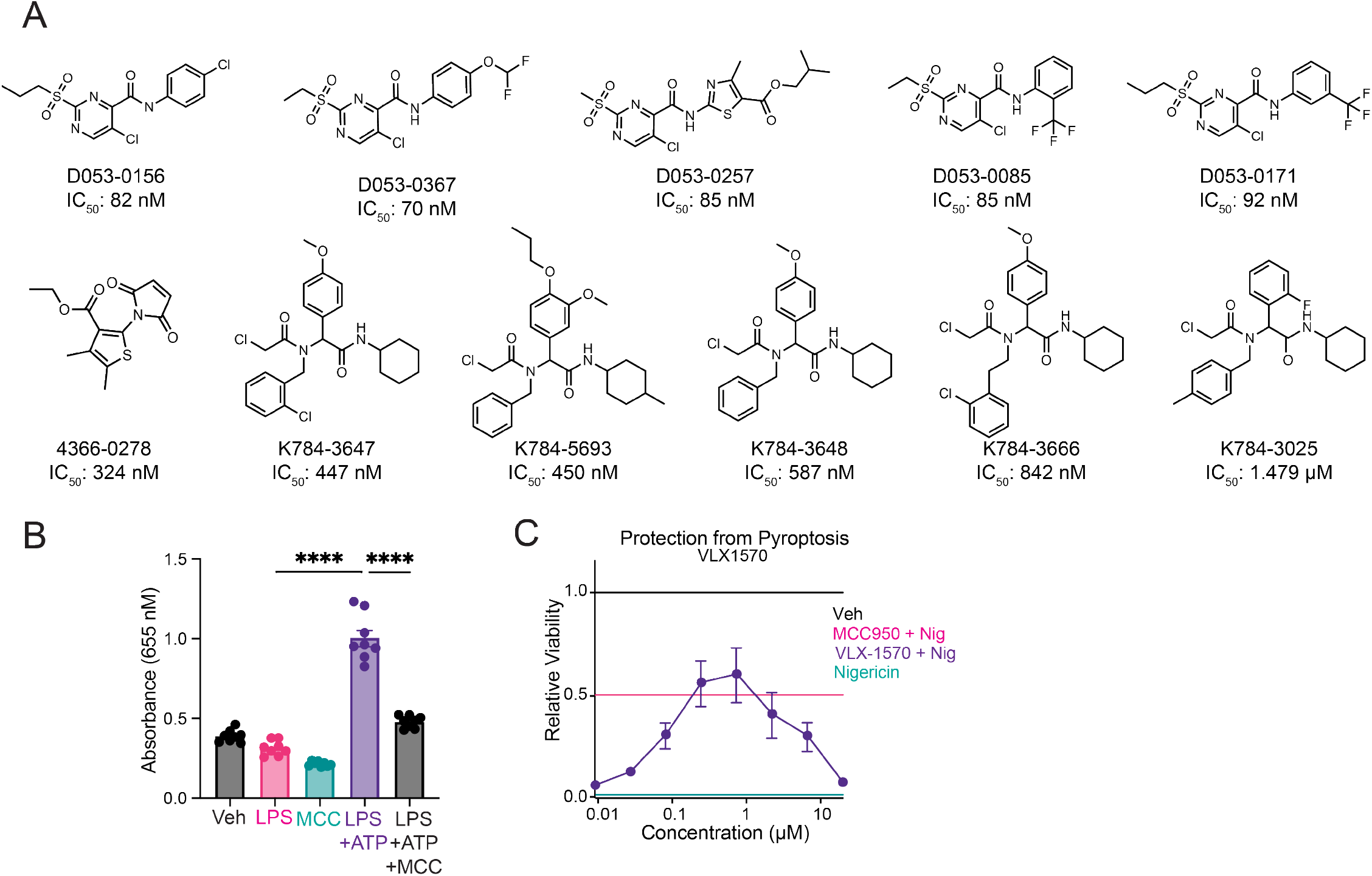
Identified hit compound classes are active in secondary assays of NLRP3 inflammasome activity. (A) Structures representing hit compounds profiled from the 9 covalent scaffolds identified. (B) IL-1β secretion inhibition in WT THP-1 cells stimulated with LPS and ATP and measured by SEAP secretion from HEK-Blue-IL-1β reporter cells. Error bars show SEM for n = 8 replicates. ****p<0.0001 for ordinary one-way analysis of variance (ANOVA) with Tukey correction for multiple comparisons between conditions. (C) Representative protection from NLRP3-mediated pyroptotic cell death induced by LPS and Nigericin in WT THP-1 by VLX1570 in dose response. Error bars show SEM for n = 4 replicates.

**Supplementary Figure 3.**
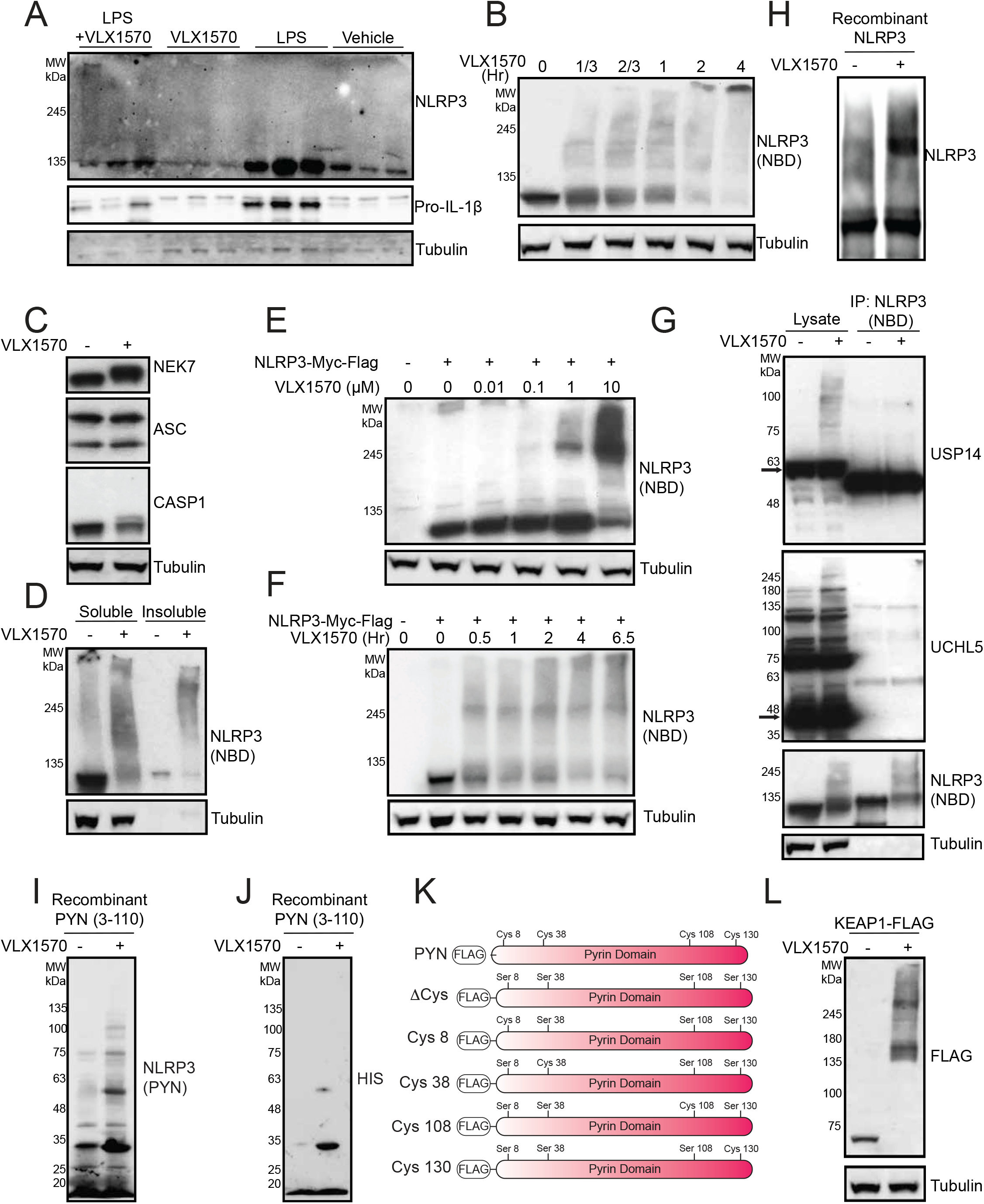
VLX1570 covalently modifies and crosslinks NLRP3. (A) Western blot of NLRP3, Pro-IL-1β, and Tubulin in cells harvested from mouse peritoneal lavage fluid of mice stimulated with 10 mg/kg LPS (3 hr), pretreated with 4.4 mg/kg VLX1570 1 hr prior to LPS injection. (B) Western blot of NLRP3 (NBD) and Tubulin from THP-1 cells treated with 10 μM VLX1570 at indicated time points. (C) Western blot for NEK7, ASC, CASP1 and Tubulin in THP-1 cells treated with 10 μM VLX1570 for 2 hr. (D) Western blot for NLRP3 (NBD) and Tubulin in soluble and insoluble fractions from lysates of WT THP-1 cells treated with 10 μM VLX1570 for 2 hr. (E) Western blot of NLRP3 (NBD) from HEK293T cells overexpressing NLRP3-Myc-FLAG treated in dose response with VLX1570 for 2 hr. (F) Western blot of NLRP3 (NBD) from HEK293T cells overexpressing NLRP3-Myc-FLAG treated in with 10 μM VLX1570 at indicated time points. (G) Western blot of USP14, UCHL5, NLRP3 (NBD), and Tubulin from WT THP-1 lysate or NLRP3-immunoprecipated material from WT THP-1 cells with or without VLX1570 treatment. (H) Western blot of NLRP3 (NBD) from recombinant WT NLRP3 treated with 50 μM VLX1570 at 4 °C overnight. (I-J) Western blot of NLRP3 (PYN) and HIS-TAG from recombinant NLRP3 PYN (3-110, C08S, C38S) treated with 50 μM VLX1570 at 4 °C overnight. (K) NLRP3 PYN Domain ΔCYS or Single CYS constructs. (L) Western blot of FLAG from HEK293T cells overexpressing KEAP1-FLAG treated with or without VLX1570 (10 μM) for 2 hr.

**Supplementary Figure 4:**
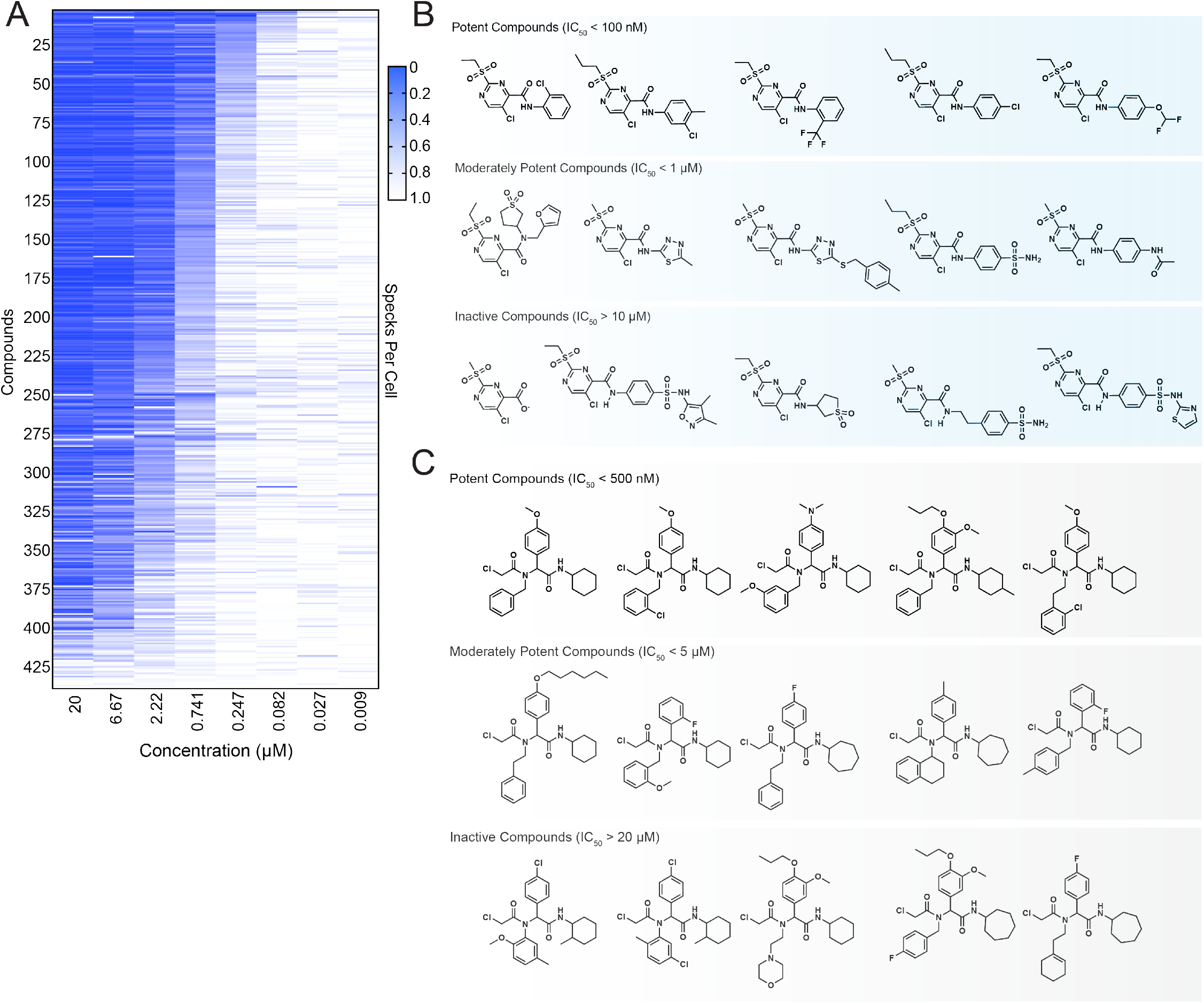
Structure activity relationship profiling of 2-sulfonylpyrimidine and chloroacetamide scaffolds. (A) Structure activity relationship for ASC-Speck Assay for chloroacetamide scaffold by supplier inventory. (B) Structure of 2-sulfonylpyrimidine compounds which are potent (IC_50_ < 100 nM), moderately active (IC_50_ < 1 μM), or inactive (IC_50_ > 10 μM) in the ASC-Speck Assay. (C) Structure of chloroacetamide compounds which are potent (IC_50_ < 500 nM), moderately active (IC_50_ < 5 μM), or inactive (IC_50_ > 20 μM) in the ASC-Speck Assay.

**Supplementary Figure 5:**
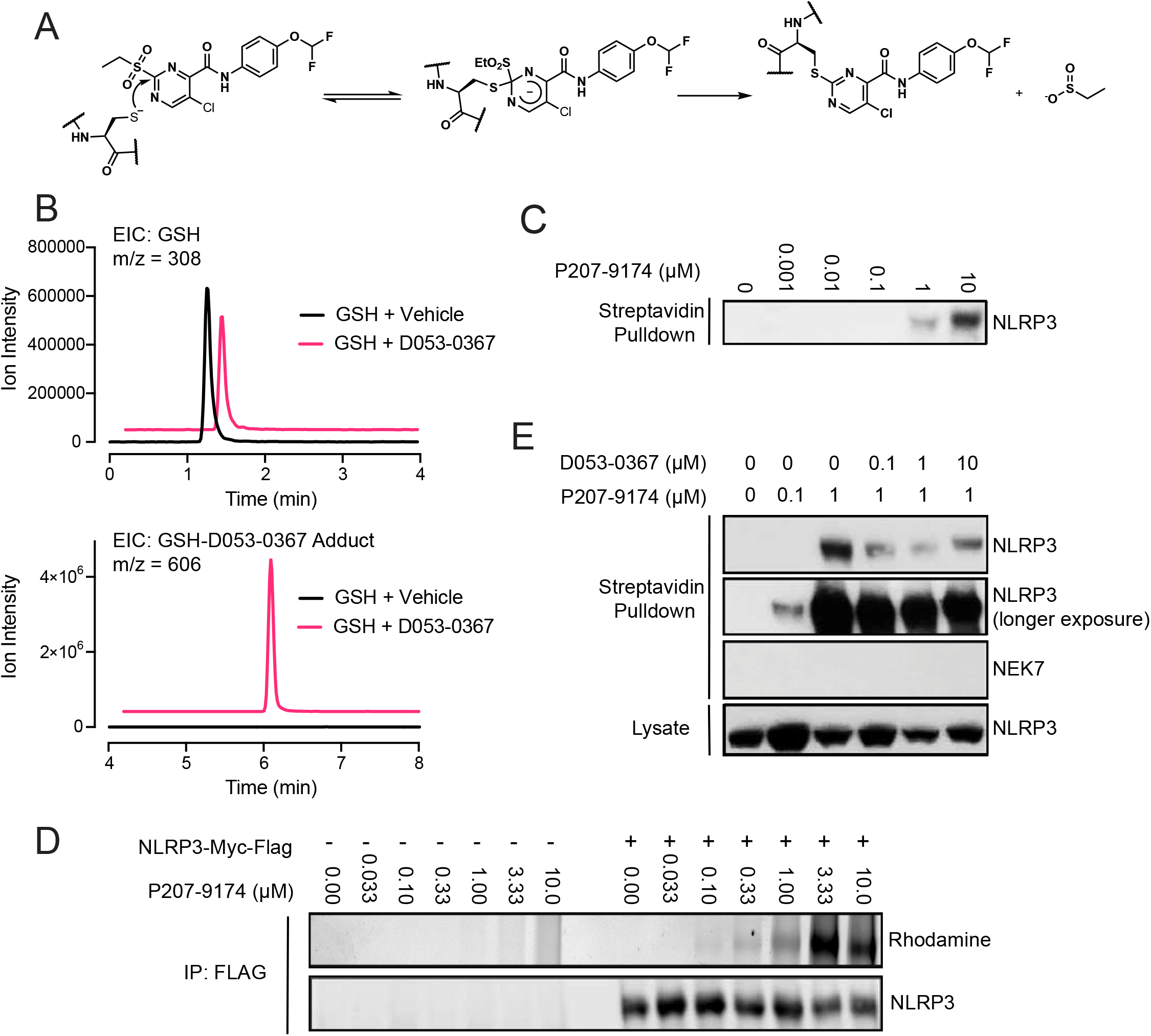
A 2-sulfonylpyrimidine-containing inhibitor of NLRP3 inflammasome formation that covalently modifies cysteines via an S_N_Ar-based mechanism. (A) S_N_Ar Covalent mechanism for 2-sulfonylpyrimidine scaffold. (B) LC-MS quantification of GSH (4 mM), D053-0367 (1 mM), or D053-0367 (1 mM) incubated in the presence or absence of GSH (4 mM) in PBS for 1 hour. Relative ion intensities were compared with chromatograms shown. (C) Anti-NLRP3 western blot of streptavidin-enriched material after in situ treatment of THP-1 cells with P207-9174 in dose response for 1 hr. (D) Anti-NLRP3 Western blot and rhodamine imaging of FLAG immunoprecipitated material after in situ treatment of NLRP3-overexpressing HEK293T cells with P207-9174 in dose response for 1 hr. (E) Anti-NLRP3 and Anti-NEK7 Western blot of streptavidin-enriched material after in situ treatment of THP-1 cells with P207-9174 for 1 hr competed with D053-0367 pretreated for 1 hr.

**Supplementary Table 1:**
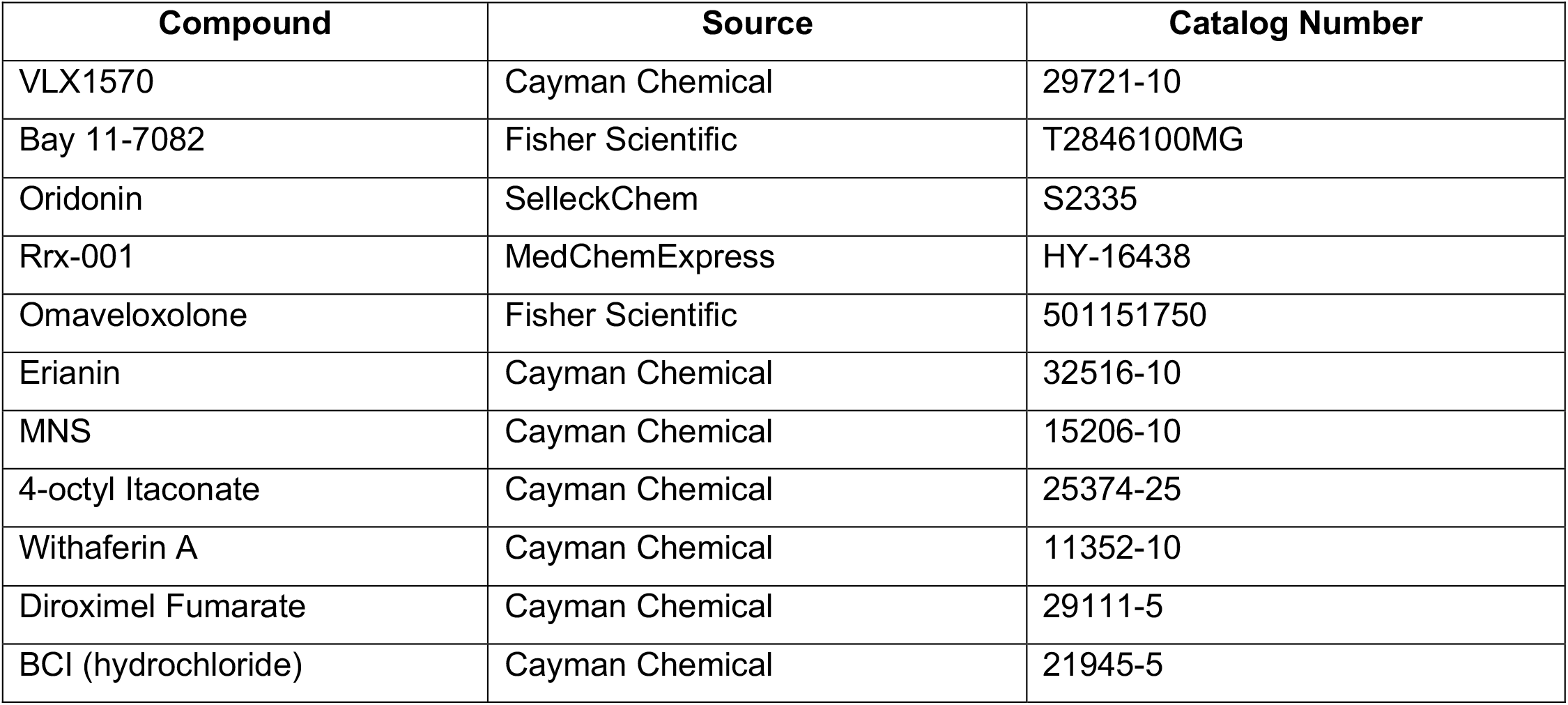

| Compound | Source | Catalog Number |
| --- | --- | --- |
| VLX1570 | Cayman Chemical | 29721-10 |
| Bay 11-7082 | Fisher Scientific | T2846100MG |
| Oridonin | SelleckChem | S2335 |
| Rrx-001 | MedChemExpress | HY-16438 |
| Omaveloxolone | Fisher Scientific | 501151750 |
| Erianin | Cayman Chemical | 32516-10 |
| MNS | Cayman Chemical | 15206-10 |
| 4-octyl Itaconate | Cayman Chemical | 25374-25 |
| Withaferin A | Cayman Chemical | 11352-10 |
| Diroximel Fumarate | Cayman Chemical | 29111-5 |
| BCI (hydrochloride) | Cayman Chemical | 21945-5 |

**Supplementary Table 2:**

| Protein | Supplier | Catalog Number | Host Species | Dilution |
| --- | --- | --- | --- | --- |
| NLRP3 (NBD) | Cell Signaling | 15101S | Rabbit | 1:1000 (BSA) |
| NLRP3 (PYN) | Adipogen | AG-20B-0014-C100 | Mouse | 1:1000 (BSA) |
| Tubulin | Sigma | T6557 | Mouse | 1:2000 (BSA) |
| FLAG | Sigma | F1804 | Mouse | 1:1000 (Milk) |
| IL-1β | GeneTex | GTX74034 | Rabbit | 1:1000 (BSA) |
| GFP | Abcam | Ab290 | Rabbit | 1:1000 (BSA) |
| USP14 | Cell Signaling | 11931S | Rabbit | 1:1000 (BSA) |
| UCHL5 | Santa Cruz | sc-271002 | Mouse | 1:1000 (BSA) |
| ASC | Cell Signaling | 13833S | Rabbit | 1:1000 (BSA) |
| CASP1 | Cell Signaling | 3866S | Rabbit | 1:1000 (BSA) |
| NEK7 | Cell Signaling | 3057S | Rabbit | 1:1000 (BSA) |

